# Metagenomic Analysis Reveals the Transfer of Antibiotic Resistance from Manure-Fertilized Soil to Carrot Microbiome

**DOI:** 10.1101/2023.11.01.565077

**Authors:** Qing Yang, Kaiqiang Yu, Jianshi Huang, Edward Topp, Yu Xia

## Abstract

Crops grown in manure-fertilized soil are more likely to be contaminated with antibiotic resistance genes (ARGs) than when grown in unmanured soil, and therefore more likely to represent a route of ARG exposure to consumers. Existing studies scarcely focused on the residual ARGs on the carrot peel growing in the manured soil after the careful washing. In the present study, residual microbiome and resistome on the carrots surface (*Daucus carota* subsp. *sativus*) at harvest after careful washing was investigated to reveal the impact of manure fertilization. Residual surface bacteria were recovered from the peel, and total bacterial DNA was extracted for the high-throughput sequencing-based metagenomic analysis. Despite the overall unaltered level of alpha diversity in both soil and carrot peel samples, manuring increased the resistome in soil significantly, but not on the carrot peel. The largely overlapped resistome detected on carrots grown with and without manuring, plus the pattern revealed by source tracking analysis indicated a soil-source of ARGs on carrots, whereas the beta-lactamases *CTX-M-84*, *OXY-4*, and *CTX-M-122* only detected in the manured soil and on the carrot peel harvested from manured soil indicated that beta-lactamases appear to be transferred from manure to the carrot. The evident impact of soil resistome and community on carrot peel microbiome, plus the limited level of plasmid and integron mediated ARGs transfer suggested the main ARGs transfer pathway from manured soil to carrot peel was via the colonization of rhizosphere soil microbes. To further elucidate the ARG propagation within the soil-carrot system, a network was constructed to explore the pattern of nine types of concurrent ARGs genotypes carried by ten different host populations, emphasizing that the residual antibiotic resistance’s transfer via raw carrot represents a risk to human health even after extensive washing.

## Introduction

Antibiotics are used extensively in human medicine and in food animal production systems for the treatment and prevention of bacterial infections. Selection pressure due to their widespread use has given rise to antibiotic resistant bacteria. The increasing difficulty in successfully treating bacterial infections now represents an urgent public health challenge that must be met with concerted global action (Vanderhaeghen and Dewulf 2017; McEwen and Collignon 2018;Tiedje et al. 2019).

A crucially important agricultural practice is the recycling of nutrients and organic matter excreted by food-producing animals into crop production systems(Hatfield and Sauer 2020). This reduces the need for chemical fertilizers, particularly important in lower income settings. However, antibiotic resistant bacteria enriched in antibiotic-treated animals, as well as intact antibiotic chemical residues, will be entrained into crop production systems with the application of the manure(Zhu et al. 2013; Sarmah et al. 2006). A number of studies have reported that growing vegetables in soils fertilized with animal manure increases the likelihood that they will carry more antibiotic resistance genes (ARGs) relative to vegetables grown in the absence of manure (Zhu et al. 2017; ; Zhang, Hu, Chen, Singh, et al. 2019).

Typically, antibiotic resistance genes were isolated from environmental samples by cloning from cultured bacteria or by PCR amplification(Riesenfeld et al. 2004). Those methods ignored potential antibiotic resistance reservoirs because most bacteria are not culturable, and PCR detection depends on primers that are based on known resistance genes. The development of culture-independent techniques was required to identify novel resistance genes and access the genetic diversity of most bacteria. High-throughput sequencing (HTS)-based metagenomic analysis represents the most robust approach to provide an integrative picture of the environmental resistome (Schmieder and Edwards 2012). It has been used to reveal the broad-spectrum profile of ARGs in various environmental matrices (Li et al. 2020; Zhao et al. 2018; Xia et al. 2017). The ARG pattern in the soil and vegetables have been explored by metagenomic technology in some studies (Xiao et al. 2016;Wind et al. 2021, Fan et al. 2021), for example, Fan et al. (Fan et al. 2021) profiled tetracycline-resistant genes and MGEs to identify the potential transmission pathways by metagenomic analysis in the soil-plant system. Wind et al. (Wind et al. 2021) analyzed dairy-derived manure and compost amendment samples, soil samples, and lettuce samples via shotgun metagenomics to assess multiple pre-harvest factors that determine lettuce resistomes. The study demonstrated a comprehensive approach to identifying key control points for the propagation of ARGs in vegetable production systems, identifying potential ARG-MGE combinations that could inform future surveillance, showing that even with imposing manure management and post-amendment wait periods in agricultural systems, ARGs originating from manure can still be found on crop surfaces. Nevertheless, few studies focused on the residual antibiotic resistomes of the plants grown on the manure-fertilized soil even after careful washing, whereas systematic evaluation of the main ARGs transfer pathway from manured soil to the harvested vegetables were scarce as well. Additionally, horizontal gene transfer (HGT) between soil resistome and carrot resistome also plays an important role in ARGs propagation. Soil bacteria, e.g. *Proteobacteria*, could facilitate the HGT of ARGs-carrying conjugative plasmids from soil into the plant endophytes (Xu et al. 2021). However, the overall transmission machinery enabled by different types of mobile vehicles were yet to be explored in manure-soil-plant system.

Taken together, the HTS-based metagenomic analysis was used in this study to investigate the ARGs profile and carrier populations to reveal exchange of antibiotic resistance between manure-fertilized soil and carrot microbiomes. In order to evaluate the potential impact of fecal material used as fertilizer, carrots were grown in soil fertilized with swine manure at agronomic rates of application. By employing the HTS-based metagenomics targeting ARGs subtypes against all the major categories of antibiotics, we aimed to (i) investigate the different influences of residual antibiotic resistance that manuring has exerted on the carrot and soil resistome; (ii) explore the potential antibiotic resistance transfer path in the carrot-soil system. The results presented will provide detailed genomic insight into the residual antibiotic resistance on edible carrot after careful washing thereby providing useful data for accessing the human resistance exposure in daily life.

## Materials and Methods

### Sampling, DNA extraction and metagenomic sequencing

#### Field operations

Experiment (E31-17-11) was undertaken during the 2017 growing season on the Agriculture and Agri-Food Canada research farm in London, Ontario, Canada (42.984°N, 81.248°W).

### Swine manure application

The application of liquid swine manure was applied at 8600 usg/ac. (79,475l/ha) on Day 136. The liquid application rate was based on manure content of crop-available nitrogen determined with an AgrosNquick test meter (Agros, Lidköping, Sweden). Manure was obtained from a local hog finisher. Finishing 1000 hogs all in/out. Manure is stored in a 250,000 gallon under barn pit. Manure was applied using a Nuhn tanker (Sebringville ON). Based on soil samples and past cropping history, there was no application of inorganic fertilizer in the control or swine plots prior to planting the plots. Immediately following application, manure was soil incorporated to a depth of 15 cm using a Kongskilde “S” tine cultivator (Denmark). Manure was applied to plot area 8mx85m separated by 3 meter borders from surrounding plots. The control plot area was 8mx85m. The layout of the farm caused this difference. Pre-planting tillage was performed on Day 185 for weed control to a depth of 15 cm using a Kuhn rototiller (Kuhn N.A. Inc, Broadhead Wisc.) and followed by a Kongskilde “S” tine cultivator (Denmark). Carrots were planted on Day 185 and grown in four 4x6m sub-plots within the 8x85m plot area. Carrot Variety (Daucus carota) Ibiza hybrid; 30cm row spacing and thinned at emergence. Plots were packed to conserve moisture with a crowfoot packer (RJ Equipment, Blenheim, ON). Insect pressures were monitored, but no action was necessary. Weed pressure in the growing season was monitored, and weeds were removed by hand hoeing and at times mechanical cultivator.

### Soil sampling

Soil cores were taken haphazardly throughout the study period, initially at pre-application and on days 0, 7, and 30, and then at crop harvest date. Six 2-cm wide cores were sampled from each of replicated carrot plots to a depth of 15 cm using a T-sampler rinsed with 70% ethanol between samplings. Cores were bulked into a labeled Ziploc bag, mixed by hand until homogenous, and transported to the laboratory in a cooler with cool packs. Thus, there were two independent soil samples analyzed at each sampling time from control and swine plots.

### Harvest

Because of the size and scope of the project, harvest occurred over four days, consisting of manured and control plots, on Days 325, 331, 333 and 338. Hand digging 10m of carrots from control and manured plots. Soil core to 15cm taken at the same time following the same above protocol. The samples of carrots and soil were delivered to the lab for further processing, following the protocol below.

### Carrot Peel Processing

In header house, wearing nitrile gloves, we wiped excess soil from carrot with dry J-cloth. We used a new J-cloth between control and treated. Wearing nitrile gloves, we carefully placed vegetables into large Ziploc bag (approx. 20 carrots (10lbs)) and add 300ml of sterile MQ water. Next carrots were carefully removed from the bag and placed into a second Ziploc bag. Another 200ml of sterile MQ water was added to the bag containing the pre-washed carrots. Gently shake bag with water for 1 minute. Next, water from bag were pooled into two labelled falcon tubes for DNA extraction.

### Soil Processing

6 cores from each plot were picked with EtOH rinsed sampler and placed in labelled plastic Ziploc bag. The soil cores were mixed homogeneously before taking 50g of soil into sterile filter stomacher bag for DNA extraction.

In summary, two carrot peel wash and two soil DNA samples from vegetables grown in ground without application of manure, or with swine manure were chosen for the subsequent DNA extraction and next-generation sequencing, namely: natural soil without manure fertilization (control-soil); fertilized soil (treated-soil); natural carrot peel grown in the natural soil (control-peel) and carrot peel grown in the manure fertilized soil (treated-peel).

### DNA extraction and Hiseq sequencing

DNA was extracted from the soil and peel-wash sample using the DNeasy PowerSoil kit (Qiagen, Canada) with default protocols. The extracted DNA was quality checked by NanoDrop ND1000 microspectrophotometer (NanoDrop Technologies, Wilmington, DE) before subject to Illumina sequencing with PE150 strategy on Hiseq 4000 platform at The Centre for Applied Genomics (The Hospital for Sick Children, Toronto, ON) (detailed procedure for sequencing library construction were shown in the supplementary).

### QC and rarefaction of Illumina dataset

To ensure all reads contained Q score >20 high-quality bases, raw reads were quality controlled by Prinseq (reference). The size of datasets is summarized in Table S1. Rarefaction curves were used as a qualitative method to estimate the ARG and taxonomic alpha diversity and richness as a function of sequencing depth. Rarefaction analysis of ARGs and bacterial communities were conducted per 5Gbp of raw reads.

### ARG quantification based on Illumina short reads

ARG abundance was determined using ARGs-OAP, which showed > 99% accuracy in ARG identification (Yin et al. 2018). Briefly, potential ARG-like reads were extracted based on Diamond against the SARG database (April 2021 version). The SARG database identifies ARG types (against specific antibiotic class) and subtypes (genotypes of ARGs within the type). Reads were regarded as valid ARGs only when they showed E value > 10-7, sequence identity > 90% as well as alignment length more than 25 amino acids to a known ARG reference. ARG abundance was normalized against cell number estimated based on Essential Single Copy Genes (ESCGs) and expressed as copies of ARGs per cell. The hierarchy structure of the SARG database allowed the ARG types and subtypes to be screened concurrently.

### Metagenome assembly

Post-QC clean reads were *de novo* assembled using the CLC Genomics Workbench (version 12.0.1, QIAGEN Bioinformatics, Denmark) with default parameters. Only contigs longer than 1kb were kept for subsequent analysis. Next, reads were mapped back to the assembled contigs using Bowtie2 (with --sensitive option) to get coverage value for each contig for subsequent quantification.

### Taxonomic assignment of ARGs to bacterial hosts

Assembled contigs were subject to open reading frame (ORFs) prediction with Prodigal (Hyatt et al. 2010). ARGs-like ORFs were determined using hmmsearch (Eddy 2009) against the SARGfam database (April 2021 version) at E value ≤ 10^-5^ with gathering threshold cutoffs (-ga). Contigs were phylogenetically assigned by combined taxonomic annotation of taxator-tk and KRAKEN (Wood and Salzberg 2014). Phylogenetic identification of ARG host populations was determined based on the phylogenetic annotation of contigs carrying ARG-like ORFs.

### ARG genotypes occurred in different samples

ARG genotypes occurred among four sample types were identified by firstly aligning ARGs-like ORFs obtained from different datasets against the customized SARG database (April 2021 version) with Lastal (Kiełbasa et al. 2011) using gap existence cost of 11. Next, the hits were filtered with the sequence similarity ≥ 99% over >90% of the query length to keep the identical instances. And two genotypes were regarded as shared in different samples when they showed an identical hit to the same ARG reference.

### Identification of integrons and plasmids

Platon (Schwengers et al. 2020) was used to filter the putative plasmids with default parameters. Integron_Finder (Nawrocki and Eddy 2013; Eddy 2011) was used for the identification of complete integrons, which is composed of the gene of *intl* , *attc* site and *attl* site.

### Statistics analysis and source-tracking of ARGs

Statistical comparisons of ARGs and microbial community between carrot peel and soil were done using nonparametric Kruskal-Wallis tests. A P-value of < 0.05 was regarded as statistically significant. Hierarchical clustering was conducted based on Bray-Curtis distance among samples. Principle Coordinate analysis (PCoA) and Venn analysis were implemented using RStudio with the R vegan package (Oksanen et al. 2012). Sourcetracker2 (Knights et al. 2011) was used to evaluate the contribution of different sources on the resistome of carrot peel harvested from manured-soil, with treated peel samples as the sink, while control peel sample and treated soil sample as the source.

## Results and Discussion

### Diversity and abundance of ARGs in the soil and on carrots

The ARG-OAP2 pipeline of the cleaned data illustrated that the average ARG abundance in control-peel group was 1.96E-02 ± 4.98E-02 ARG copy per cell , which was slightly higher than that of the treated-peel group of 1.72E-02 ± 4.52E-02 copy per cell. The average ARGs concentration in the control-soil was 2.53E-02 ± 6.09E-02 copy per cell , which was lower than that of the treated-soil group of 3.18E-02 ± 8.30E-02 copy per cell (Figure S1a). After *de novo* assembly, 92,036 contigs were obtained in the peel sample with an average length of 1,976bp while 74,483 contigs were obtained in the soil sample with an average length of 1,945bp. Based on HMM-based function prediction, we annotated 3,659 and 3,771 ORFs as ARGs-like ORFs of carrot peel and soil samples. The detailed information of the metagenomes can be found in Table S1.

Among 17 categories of ARG types identified, the predominant ARG types in all samples were multidrug, beta-lactam and aminoglycoside resistance genes, covering three major resistance mechanisms: antibiotic deactivation, efflux pump and cellular protection. 172 ARG subtypes were detected in all samples. 79 resistant genotypes showed the same trend of variation (46 genotypes of enrichment and 33 genotypes of depletion) between treated and control group in soil and peel samples (red dot and the corresponding blue dot on the same side of the middle line in Figure 1). These genotypes included: 47 of beta-lactam, 13 of multidrug, 3 of vancomycin, 2 of MLS, 2 of chloramphenicol, 1 of aminoglycoside, 1 of fosfomycin and 10 of others, which respectively took 11.25% and 12.8% of soil and peel resistome, indicating the overall consistent impact of manure fertilization on soil and peel resistome. Previous studies found a similar trend of drug resistance in the manured soil and lettuce grown in it (Zhang, Hu, Chen, Singh, et al. 2019). Curiously, some studies reported that *tet* genes increased significantly after manure application (Leclercq et al. 2016; Xiong et al. 2018). However, the abundance of tetracycline resistance genes found in our study, namely *tet34*, *tet35*, *tetA* and *tetV* were fairly low and the influence exerted by manuring application on them was negligible. Such inconsistent resistome alternation indicates the impact of manure application could be application-specific and affected by the combined effect of the resistance composition of manure applied and soil microbiome.

**Figure 1.**
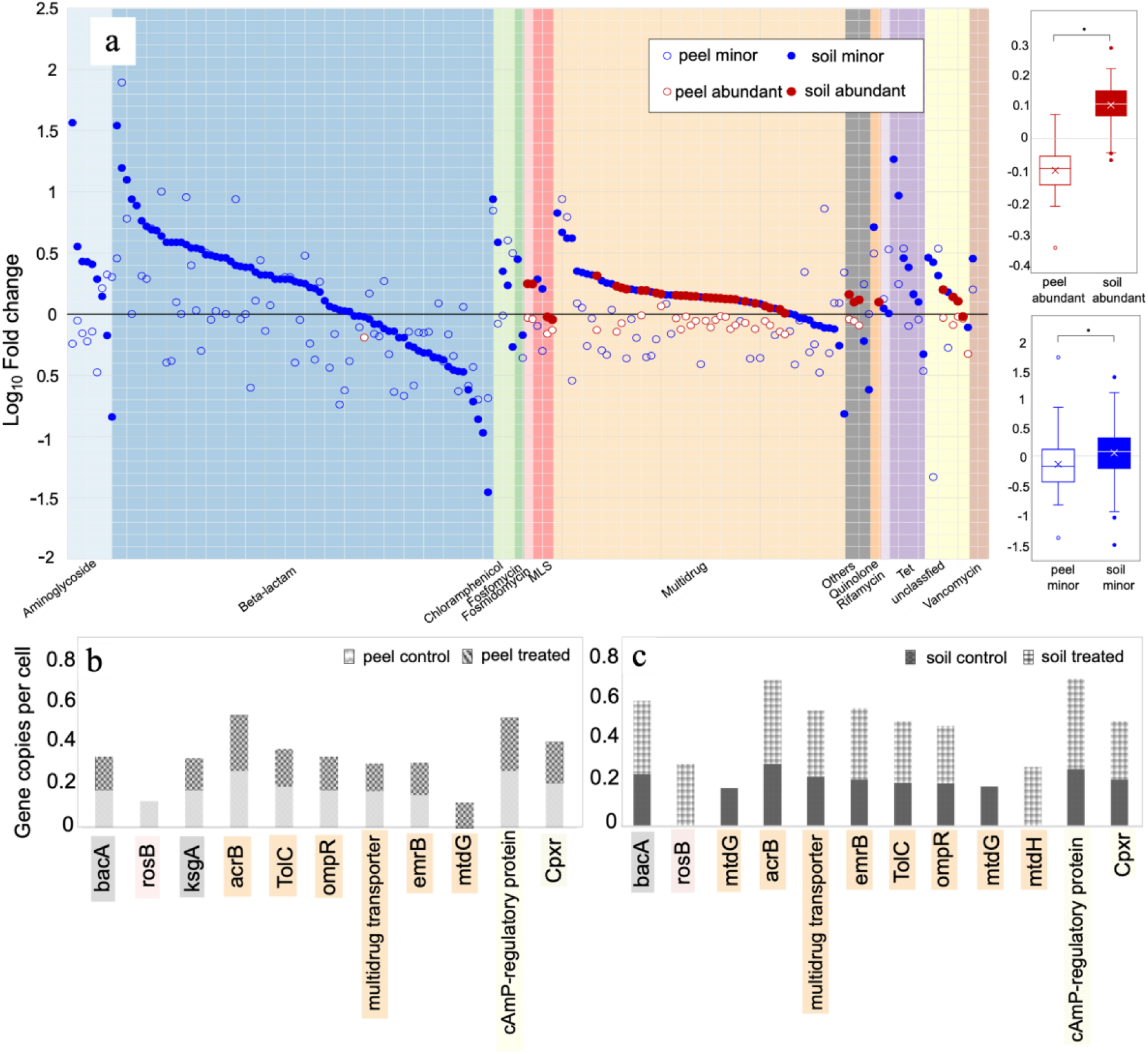
(a) The overall trend of resistome variation on the carrot peel and soil after manure fertilization. ARGs are quantified based on the metagenomic reads. If a ARG showed abundance higher than the average abundance value of the overall resistome, it will be categorized as “Abundant” in the plot, otherwise it will be considered as “Minor”. The boxplots on the right show the comparison of fold change between soil and peel samples with “Abundant” and “Minor” ARGs compared separately (Paired-sample *t-test*). (b) and (c) respectively depicts the distribution of the top10 dominating ARGs of carrot peel and soil samples.

Given that both ARGs and microbial communities were notably altered with fertilization, we sought to investigate the antimicrobial genetic resistance and microbial variation between natural and manured groups. We found that manuring increased the abundance of ARGs in soil significantly but not as evident on carrot peel. We first calculated the number of ARGs enriched (fold change >1) or depleted (fold change <1) in peel and soil with fertilization and then focused on those subtypes that are abundant (with relative abundance above the average level, shown as filled dots in Figure 1). As shown in the boxplot of Figure 1, both abundant and minor proportion of the peel resistome showed significantly (p < 0.01) lower variation than that of the soil resistome, suggesting the resistome of carrot peel samples was more recalcitrant to the influence brought by manure fertilization compared to soil samples. Such a pattern was also observed in a previous study that a larger variety of ARGs and higher detection frequency of resistant coliforms in soils than on the harvested vegetables (Marti et al. 2013). Presumably, the conservativeness of peel resistome towards manure fertilization was attributed to the cleaning step which washed off the soil from harvested carrots, as would be expected in normal culinary preparation. Besides that, a large part of high abundant ARGs tends to remain a less evident fold change after manure fertilization (Figure 1). For example, the multidrug efflux protein of *acrB* and *TolC, ABC* transporter protein (*bacA*) that regulates translocation of bacitracin, as well as several regulating factors (e.g. *CpxR*) combating a variety of extracytoplasmic protein-mediated toxicities(Danese and Silhavy 1998), all showed minimum level of variation after manure fertilization (Figure 1b and 1c). Noteworthy is that several abundant genotypes showed opposite trend of variation in soil and carrot peel resistome, for instance, *rosB*-involving in the resistance against antibiotic roseoflavin (Mora-Lugo et al. 2019), was dominant on the natural carrot peel but got depleted in treated peel, while this gene was only detected in soil with manure fertilization. Additionally, some gene targets were only detected in the manured soil and on the carrot peel when harvested from manured soil, such as the beta-lactamases *CTX-M-84, OXY-4*, and *CTX-M-122,* indicating that beta-lactamases appear to be transferred from manure to the carrot.

### Community structure of soil and carrot peel microbiome

The natural carrot peel community was dominated by *Enterobacteriaceae*, *Pseudomonadaceae,* and *Coxiellaceae* respectively taking 43.9%, 3.22% and 2.91% of the community, which was partially different to that of the soil community which showed high prevalence of *Enterobacteriaceae*, *Pasteurellaceae, Erysipelotrichaceae,* and *Comamonadaceae* (61.7%, 2.31%, 1.79%, 1.63%) (Figure S2(b)). After manure fertilization, *Prevotellaceae* and *Cytophagaceae* that took 3.98% and 3.41% outcompeting *Pseudomonadaceae* and *Coxiellaceae* became prevalent on the carrot peel, while *Pseudomonadaceae* (3.09%) replacing *Pasteurellaceae* became one of the dominant populations in manure-treated soil microbiome. Since *Enterobacteriaceae* represented the predominant population of human fecal microbiota, its high prevalence on carrot and peel highlighted the migration and installation of gut microbes in soil microbiomes. Like the ARGs profile, manuring caused stronger alteration of soil community than peel community (Figure S2). Despite the overall unchanged level of alpha diversity in both soil and peel samples as revealed by rarefaction analysis (Figure S1b,c), there were respectively 113 and 72 families that showed evident (> 1 fold) variation in peel and soil group after manure fertilization. The *Nocardioidaceae* family of *Actinobacteria* phyla held the highest increment after manure fertilization with fold change of 3.13 in the soil, while *C111* family of *Actinobacterial* phylum decreased over 100 times after the fertilization on the carrot peel.

Previous studies had shown animal manure could serve as an important reservoir of antibiotic resistant bacteria and land application of manure could introduce a number of unique ARG subtypes to agricultural field and vegetables (Chee-Sanford et al. 2009; Fernández-Alarcón, Singer, and Johnson 2011; Zhang et al. 2017; Zhu et al. 2017), but, it has not been established in the previous studies that gene targets detected more frequently on harvested vegetables were carried into the soil by manure or were carried in the local soil bacteria that got enriched by the manure fertilization. As the basis for a better understanding of this question, environmental host populations of ARGs were identified using taxonomic annotation of ARG-carrying contigs. Figure 2 indicates that assembled ARGs were harbored by diverse bacterial families including *Yersiniaceae, Enterobacteriaceae, Erwiniaceae, Pseudomonadaceae, Pectobacteriaceae, Alteromonadaceae* and etc, suggesting that the fecal resistome resided in phylogenetically diverse microbiota. *Yersiniaceae*, as the predominant bacterial host, carried a wide spectrum of ARGs including the genes resistant to multidrug, aminoglycoside, MLS, beta-lactam, vancomycin, bacitracin, quinolone, tetracycline, and polymyxin. Among the dominating hosts, peel and soil samples presented the similar components, but *Leuconostocaceae* and *Flavobacteriaceae* only showed up in the peel. The potential tendency of the ARGs transfer carried by those local communities would be explained subsequently.

**Figure 2.**
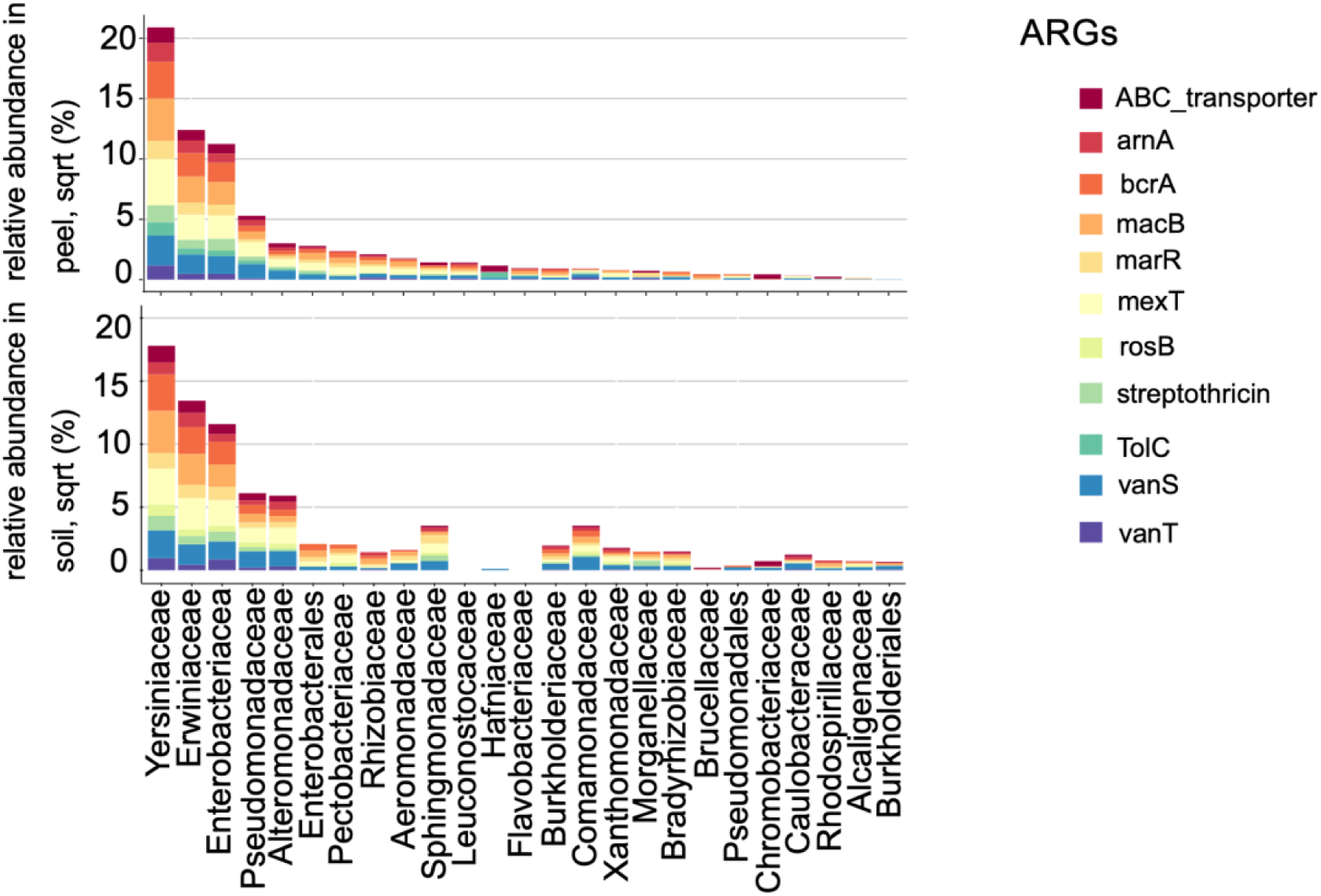
Composition of the host populations of the ten most abundant ARGs of carrot peel and soil resistome. Only dominating hosts with abundance > 0.05 % are shown in the figure.

### Human pathogenic bacteria carrying ARGs on the carrots and in the soil

We also paid special attention to ARGs genotypes with broad health threats. Although *mcr-1* and *NDM-1* were not detected, a total 14 types of *blaCTX-Ms* were detected with *CTX-M-94* dominant in both carrot peel and soil resistome based on the raw reads. Manuring enriched *CTX-M-75* by 10.0-fold on the carrot peel while *CTX-M-152* increased over 12-fold in the soil. *blacCTX-Ms* are the largest group of ESBLs (extended-spectrum beta-lactamases) enabling the resistance development of ESBL-producing *Enterobacteriaceae* (Bevan, Jones, and Hawkey 2017). Surprisingly, in the metagenomic assembly, beta-lactam resistance was no longer the dominant ARG type, especially no *blaCTX-Ms* could be detected. It is presumably explained by the fact that it is particularly difficult to correctly assemble regions containing exogenous genes acquired by horizontal gene transfer (HGT), such as resistance and prophages, owing to their inherent repetitive nature or the flanking of these genes by repetitive insertion sequences (Ashton et al. 2015), which could be best demonstrated by the widespread plasmid-mediated *blaCTX-Ms* transmission (Awosile and Agbaje 2021; Barlow et al. 2008).

Worth to note is that *Klebsiella pneumoniae* was only found on the carrot peel (81,506 and 33,099 assembly reads in the control group and treated group respectively) while it was not found in the soil. *K. pneumoniae* is an important multidrug-resistant pathogen affecting humans and a major source for hospital infections associated with high morbidity and mortality due to limited treatment options. Under antibiotic selective pressure, *K. pneumoniae* continuously accumulates ARGs, by *de novo* mutations, and via acquisition of plasmids and transferable genetic elements, leading to extremely drug resistant (XDR) strains harboring a ‘super resistome’(Campos et al. 2016). We identified 15 different ARGs genotypes in *K. pneumoniae,* highlighting the potential health threats imposed by residual pathogens on raw vegetables even after washing. Intriguingly, manuring decreased the antibiotic resistance encompassed by *K. pneumoniae* by 3-fold on the carrot peel, which may because bacteria from manure may not be well adapted to the soil environment (Heuer et al. 2011).

### The spread pathway of ARGs from soil resistome to carrot resistome

There are many potential pathways via which ARGs can transfer from manured soil to vegetables. For example, ARGs-carrying bacteria may attach to the aerial leaf surfaces of vegetables, survive as phyllosphere microorganisms, accumulating in leaf tissues as leaf endophytes (Lindow and Brandl 2003), or transfer into vegetable roots as root endophytes (Lindow and Brandl 2003; Hardoim et al. 2015). Zhang, Hu, Chen, Yan, et al. also provided evidence that enrichment of the resistome in the rhizosphere and phyllosphere is more obvious than with the endosphere after manure application. In addition, horizontal gene transfer (HGT) between soil resistome and carrot resistome also play an important role in ARGs propagation.

In this study, the potential pathway of ARGs spreading in the manure-soil-carrot system was revealed. Figure 3a showed that natural carrot peel and fertilized soil respectively contributed 48.3% and 49.9% to the fertilized carrot peel resistome, consolidating the notion that soil and manure derived ARGs are important sources of resistome found in vegetables (Q.-L. Chen et al. 2017). The importance of rhizosphere soil microbes on carrot resistome could be further evidenced by the source tracking analysis based on ARGs host that manured soil microbes contributed to 53.6% of the host community on carrot peel, which was even higher than the influence of native host populations on carrot (43.9% of the carrot control group) (Figure 3b).

**Figure 3.**
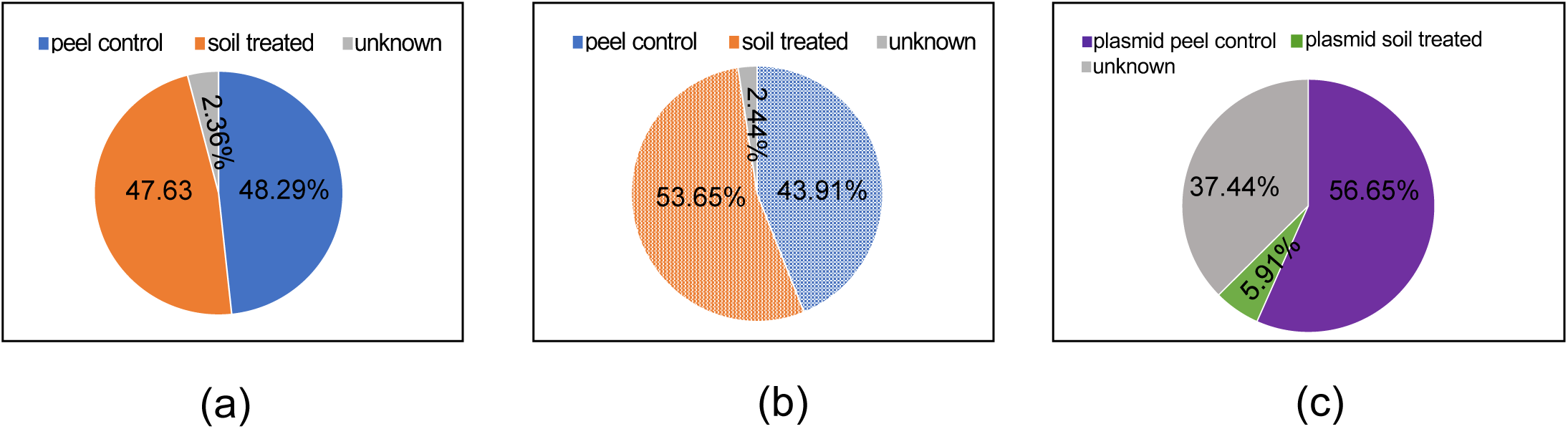
Bayesian source tracking results. (a,b) ARGs and bacterial community based on the metagenomic assembly - manured peel as the sink, control peel, manured soil as the source (c) Plasmid-mediated ARGs, which regards plasmids carrying ARGs on the manured peel as the sink, plasmid carrying ARGs on the control peel and in the manured soil as the source

Plasmids respectively took 0.391% and 0.453% (quantified based on the reads aligned to plasmid-like contigs) of carrot peel and soil metagenomes. In contrast, no ARGs-containing integron could be detected, indicating the unlikeliness of integron-mediated ARGs dissemination in our manure-soil-carrot system. As shown in Figure 4, the dominant components of plasmid resistome included ARGs types of multidrug, bacitracin, MLS as well as vancomycin. Four genotypes of *mexT, vanS, bcrA* and *macB* appeared on plasmids on the carrot peel sample, while eight genotypes were detected in the soil samples, with *mexT, vanS, qnrB, macB* and *bcrA* being the most prevalent. Two of plasmid-carried ARGs genotypes on carrot peel, namely *macB* and *bcrA*, showed at least doubled abundance increment after manure fertilization. Similar enrichment of plasmids-mediated resistomes caused by manure application was also observed in carrot tissues (Mei et al. 2021). Since plasmid-encoded ARGs on the treated peel sample is a combination of 1) plasmid ARGs carried by the native carrot peel populations and 2) plasmid ARGs transmitted from the treated soil microbes, we put the resistome of these two components as the potential “source” for source tracking analysis with plasmids carrying ARGs on treated peel as the “sink”. Compared to the native plasmid carrying resistome of carrot (the carrot control group), ARGs obtained via plasmid-mediated transmission from soil microbes (the treated soil group) showed minor contribution on the plasmid carrying ARGs on carrot peel after manure fertilization (Figure 3c). Taken together, the evident impact of soil resistome and community on carrot peel microbiome, plus the limited level of plasmids-mediated ARGs HGT, all suggested the main ARGs transfer pathway from manured soil to carrot peel is via the colonization of rhizosphere soil microbes.

**Figure 4.**
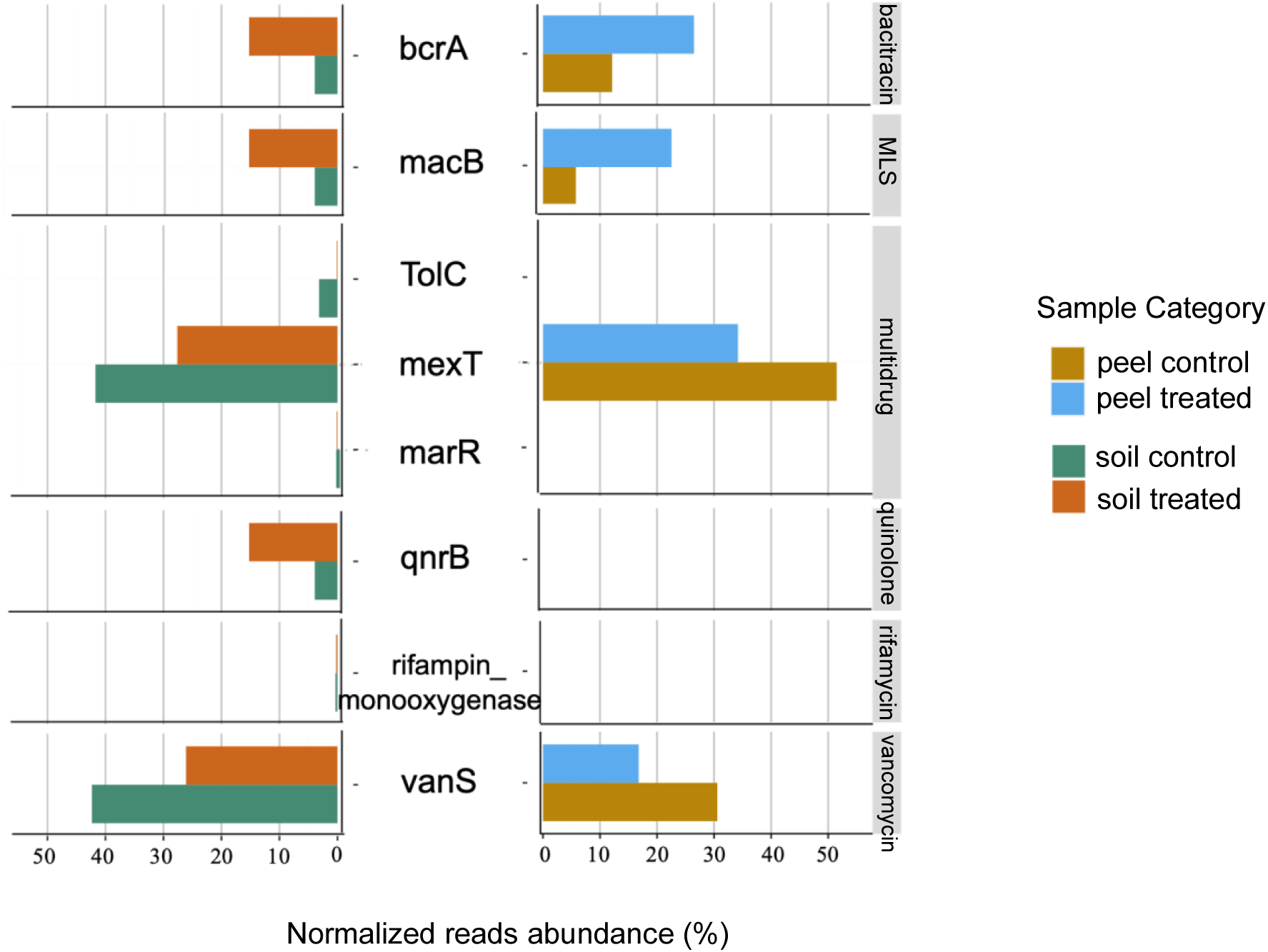
ARGs of carrot peel and soil mediated by the plasmids

To further elucidate the ARG propagation within the manure-soil-carrot system, a network was constructed to explore the pattern of concurrent ARGs genotypes among different host populations in soil and carrot peel samples with and without manure application. To ensure reliability of the genotype sharing network, a stringent cut-off (> 95% sequence similarity over >90% of the corresponding gene length) was used to identify sharing instances. Nine concurrent ARG genotypes of *marR, bcrA, arnA, ksgA, mexT, macB, ykkD, tet34* and *ykkC* appeared among carrot peel and soil samples. The abundance of these genotypes respectively increased from 41.9% to 43.5% on carrot peel and from 39.5% to 43.8% in soil after manure application. Except for *marR*, manuring stimulated the abundance of all other ARGs with *tet34* acquired the most increase around 7 times. Additionally, the relative abundance of polymyxin resistance genes (*arnA*) was much higher on the carrot peel than in the soil, whereas other resistance genotypes showed relatively similar distribution between soil and carrot peel. Worth to note is that all the shared genotypes as well as their carrying populations in the treated group (peel treated and soil treated), could be identified in the control group without fertilization (Figure 5), indicating that no exogenous sharing pattern had been introduced after fertilization, again highlighting the recalcitrance of peel resistome against manure fertilization.

**Figure 5.**
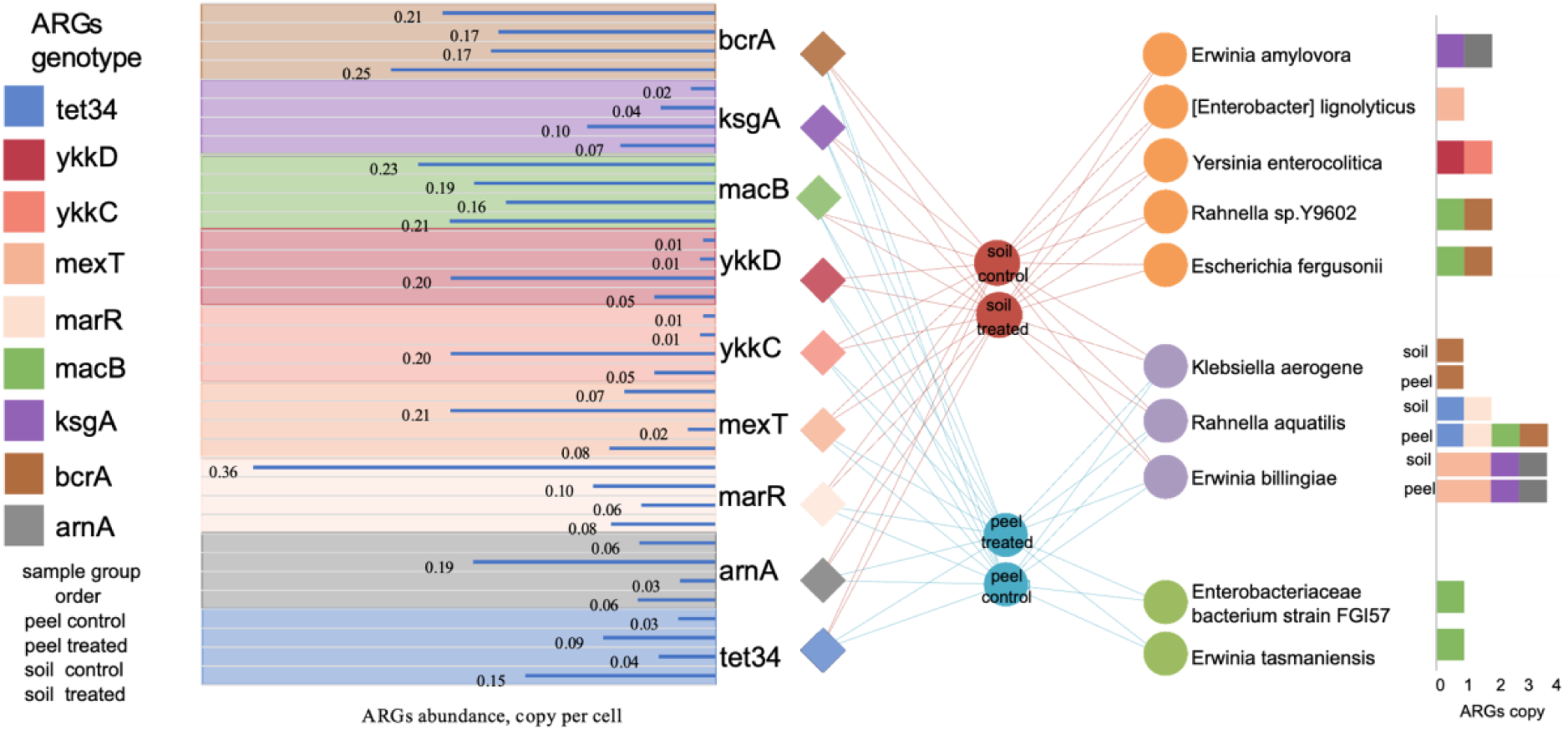
Network depicts the distribution of shared ARGs genotypes (left) and their host populations (right) among soil and peel samples. Bar chart on the left illustrates the abundance of these shared genotypes among the four sample groups studies. For each genotype, the top to bottom sequence of the four bars are peel control, peel treated, soil control, soil treated. The copy number of ARGs encoded by the host populations are shown in the bar plot on the right. For host populations appeared in both soil and peel samples (the purple group), their ARGs profile are displayed separately.

It can also be observed that bacterial species carrying the nine concurrent ARGs were the plausible vehicle for the ARGs migration from manure. Ten bacterial species were identified as the host populations of the shared ARGs genotypes. Among them, we witnessed two species of human pathogens namely, *Rahnella aquatilis* and *Yersinia enterocolitica*. *R. aquatilis* was observed in both soil and carrot peel communities while *Y. enterocolitica* only observed in soil microbiome. Interestingly, the shared ARGs were carried by different host populations community in soil and carrot peel microbiome that only three host species of *R. aquatilis*, *Erwinia billingiae* and *Klebsiella aerogenes* were observed in both soil and carrot peel community, while *Enterobacteriaceae bacterium strain FGI 57*, *Erwinia tasmaniensis* were only detected on carrot peel and *Escherichia fergusonii*, *Erwinia amylovora, Enterobacter lignolyticus*, *Yersinia enterocolitica* and *Rahnella YP602* were only found in soil. The ARGs profile of these populations showed slight differentiation resistome between soil and peel group too, for instance, *R. aquatilis* in soil carried *marR* and *tet34*, both of which got enriched after the manure treatment, while *macB* and *bcrA* were also encompassed by this population identified on carrot peel. Regarding the overall ARG profile carried by those ten species, *R. aquatilis* was the most abundant host that carried 60.0% and 35.0% of shared ARGs in the peel and soil samples, which indicates the risk of the transmission of pathogens and carrying ARGs.

## Conclusions

In summary, we demonstrated the impacts of manure fertilization on the diversity and abundance of ARGs and the associated bacterial communities in agricultural soils and on carrots by shot-gun sequencing based metagenomics. Resistome alteration (enrichment) caused by manure application was profiled with the host populations and mobility identified with state-of-the-art metagenomic approaches. We found that in the manure-soil-carrot system, the overall resistance enrichment caused by manure application was much more evident in the soil than on the carrot peel. Plasmid-mediated ARGs are also inclined to occur from manure to the soil rather than to carrot phyllosphere, and beta-lactamases appear to be transferred from manure to the carrot. Bacteria hosts that carried concurrent manure-enriched ARGs migrating in the soil-carrot system included *Yersiniaceae, Enterobacteriaceae, Erwiniaceae, Enterobacteriaceae*, and *Burkholderiaceae*. The persistence of resistant pathogens shared between carrot peel and manured soil indicated a potential route for ARGs to transfer into human microbiome and pathogens via the food chain (e.g. vegetable, salad). With such, we concluded that antibiotic resistance transfer via raw carrot represents a risk to human health even after careful washing; therefore peeling is needed to ensure the safety usage of organically grown carrots. And in agricultural production, it is necessary to take appropriate agricultural measures to reduce the colonization of soil ARBs to roots, such as the use of rhizosphere beneficial microorganisms (e.g., nitrogen-fixing bacteria).

## Acknowledgement

The authors are grateful to the National Key R&D Program of China (Grant No: 2022YFE0103200), National Natural Science Foundation of China (42177357) and XX for financial support. Also, we want to thank the State Environmental Protection Key Laboratory of Integrated Surface Water-Groundwater Pollution Control, Center for Computational Science and Engineering at Southern University of Science and Technology (SUSTech), and core research facilities at SUSTech to provide quality resources and services.

## Research data for this article

All the sequence reads were deposited into the China National Genebank (CNGB) under the project accession number: CNP0002124

## References

1. Ashton, Philip M., Satheesh Nair, Tim Dallman, Salvatore Rubino, Wolfgang Rabsch, Solomon Mwaigwisya, John Wain, and Justin O’Grady. 2015. “MinION Nanopore Sequencing Identifies the Position and Structure of a Bacterial Antibiotic Resistance Island.” Nature Biotechnology 33 (3): 296–300.

2. Awosile, Babafela B., and Michael Agbaje. 2021. “Genetic Environments of Plasmid-Mediated blaCTXM-15 Beta-Lactamase Gene in Enterobacteriaceae from Africa.” Microbiology Research. 10.3390/microbiolres12020026.

3. Barlow, Miriam, Rebecca A. Reik, Stephen D. Jacobs, Mónica Medina, Matthew P. Meyer, John E. McGowan Jr, and Fred C. Tenover. 2008. “High Rate of Mobilization for blaCTX-Ms.” Emerging Infectious Diseases 14 (3): 423–28.

4. Bevan, Edward R., Annie M. Jones, and Peter M. Hawkey. 2017. “Global Epidemiology of CTX-M β-Lactamases: Temporal and Geographical Shifts in Genotype.” Journal of Antimicrobial Chemotherapy. 10.1093/jac/dkx146.

5. Campos, Anaelís C., James Albiero, Alessandra B. Ecker, Cristina M. Kuroda, Lívia E. F. Meirelles, Angelita Polato, Maria C. B. Tognim, Márcia A. Wingeter, and Jorge J. V. Teixeira. 2016. “Outbreak of Klebsiella Pneumoniae Carbapenemase-Producing K Pneumoniae: A Systematic Review.” American Journal of Infection Control 44 (11): 1374–80.

6. Chee-Sanford, Joanne C., Roderick I. Mackie, Satoshi Koike, Ivan G. Krapac, Yu-Feng Lin, Anthony C. Yannarell, Scott Maxwell, and Rustam I. Aminov. 2009. “Fate and Transport of Antibiotic Residues and Antibiotic Resistance Genes Following Land Application of Manure Waste.” Journal of Environmental Quality 38 (3): 1086–1108.

7. Chen, Qing-Lin, Xin-Li An, Yong-Guan Zhu, Jian-Qiang Su, Michael R. Gillings, Zhi-Long Ye, and Li Cui. 2017. “Application of Struvite Alters the Antibiotic Resistome in Soil, Rhizosphere, and Phyllosphere.” Environmental Science & Technology 51 (14): 8149–57.

8. Danese, P. N., and T. J. Silhavy. 1998. “CpxP, a Stress-Combative Member of the Cpx Regulon.” Journal of Bacteriology 180 (4): 831–39.

9. Eddy, Sean R. 2009. “A New Generation of Homology Search Tools Based on Probabilistic Inference.” Genome Informatics. International Conference on Genome Informatics 23 (1): 205–11.

10. Eddy, Sean R. 2011. “Accelerated Profile HMM Searches.” PLoS Computational Biology 7 (10): e1002195.

11. Fang, Hua, Huifang Wang, Lin Cai, and Yunlong Yu. 2015. “Prevalence of Antibiotic Resistance Genes and Bacterial Pathogens in Long-Term Manured Greenhouse Soils as Revealed by Metagenomic Survey.” Environmental Science & Technology 49 (2): 1095–1104.

12. Fan, Haonan, Shanghua Wu, Wenxu Dong, Xianglong Li, Yuzhu Dong, Shijie Wang, Yong-Guan Zhu, and Xuliang Zhuang. 2021. “Characterization of Tetracycline-Resistant Microbiome in Soil-Plant Systems by Combination of H218O-Based DNA-Stable Isotope Probing and Metagenomics.” Journal of Hazardous Materials 420 (July): 126440.

13. Fernández-Alarcón, Claudia, Randall S. Singer, and Timothy J. Johnson. 2011. “Comparative Genomics of Multidrug Resistance-Encoding IncA/C Plasmids from Commensal and Pathogenic Escherichia Coli from Multiple Animal Sources.” PloS One 6 (8): e23415.

14. Hardoim, Pablo R., Leonard S. van Overbeek, Gabriele Berg, Anna Maria Pirttilä, Stéphane Compant, Andrea Campisano, Matthias Döring, and Angela Sessitsch. 2015. “The Hidden World within Plants: Ecological and Evolutionary Considerations for Defining Functioning of Microbial Endophytes.” Microbiology and Molecular Biology Reviews: MMBR 79 (3): 293–320.

15. Hatfield, Jerry L., and Thomas J. Sauer. 2020. Soil Management: Building a Stable Base for Agriculture. John Wiley & Sons.

16. Heuer, Holger, Heike Schmitt, and Kornelia Smalla. 2011. “Antibiotic Resistance Gene Spread due to Manure Application on Agricultural Fields.” Current Opinion in Microbiology 14 (3): 236–43.

17. Hyatt, Doug, Gwo-Liang Chen, Philip F. Locascio, Miriam L. Land, Frank W. Larimer, and Loren J. Hauser. 2010. “Prodigal: Prokaryotic Gene Recognition and Translation Initiation Site Identification.” BMC Bioinformatics 11 (March): 119.

18. Kiełbasa, Szymon M., Raymond Wan, Kengo Sato, Paul Horton, and Martin C. Frith. 2011. “Adaptive Seeds Tame Genomic Sequence Comparison.” Genome Research 21 (3): 487–93.

19. Knights, Dan, Justin Kuczynski, Emily S. Charlson, Jesse Zaneveld, Michael C. Mozer, Ronald G. Collman, Frederic D. Bushman, Rob Knight, and Scott T. Kelley. 2011. “Bayesian Community-Wide Culture-Independent Microbial Source Tracking.” Nature Methods 8 (9): 761–63.

20. Leclercq, Sébastien Olivier, Chao Wang, Yaxin Zhu, Hai Wu, Xiaochen Du, Zhipei Liu, and Jie Feng. 2016. “Diversity of the Tetracycline Mobilome within a Chinese Pig Manure Sample.” Applied and Environmental Microbiology 82 (21): 6454–62.

21. Lindow, Steven E., and Maria T. Brandl. 2003. “Microbiology of the Phyllosphere.” Applied and Environmental Microbiology. 10.1128/aem.69.4.1875-1883.2003.

22. Li, Yiming, Weiwei Cao, Shuli Liang, Shinji Yamasaki, Xun Chen, Lei Shi, and Lei Ye. 2020. “Metagenomic Characterization of Bacterial Community and Antibiotic Resistance Genes in Representative Ready-to-Eat Food in Southern China.” Scientific Reports 10 (1): 15175.

23. McEwen, Scott A., and Peter J. Collignon. 2018. “Antimicrobial Resistance: A One Health Perspective.” Microbiology Spectrum 6 (2). 10.1128/microbiolspec.ARBA-0009-2017.

24. Mei, Zhi, Leilei Xiang, Fang Wang, Min Xu, Yuhao Fu, Ziquan Wang, Syed A. Hashsham, Xin Jiang, and James M. Tiedje. 2021. “Bioaccumulation of Manure-Borne Antibiotic Resistance Genes in Carrot and Its Exposure Assessment.” Environment International 157 (August): 106830.

25. Mora-Lugo, Rodrigo, Julian Stegmüller, and Matthias Mack. 2019. “Metabolic Engineering of Roseoflavin-Overproducing Microorganisms.” Microbial Cell Factories 18 (1): 146.

26. Nawrocki, Eric P., and Sean R. Eddy. 2013. “Infernal 1.1: 100-Fold Faster RNA Homology Searches.” Bioinformatics 29 (22): 2933–35.

27. Rahube, Teddie O., Romain Marti, Andrew Scott, Yuan-Ching Tien, Roger Murray, Lyne Sabourin, Yun Zhang, Peter Duenk, David R. Lapen, and Edward Topp. 2014. “Impact of Fertilizing with Raw or Anaerobically Digested Sewage Sludge on the Abundance of Antibiotic-Resistant Coliforms, Antibiotic Resistance Genes, and Pathogenic Bacteria in Soil and on Vegetables at Harvest.” Applied and Environmental Microbiology. 10.1128/aem.02389-14.

28. Sarmah, Ajit K., Michael T. Meyer, and Alistair B. A. Boxall. 2006. “A Global Perspective on the Use, Sales, Exposure Pathways, Occurrence, Fate and Effects of Veterinary Antibiotics (VAs) in the Environment.” Chemosphere 65 (5): 725–59.

29. Schmieder, Robert, and Robert Edwards. 2012. “Insights into Antibiotic Resistance through Metagenomic Approaches.” Future Microbiology 7 (1): 73–89.

30. Schwengers, Oliver, Patrick Barth, Linda Falgenhauer, Torsten Hain, Trinad Chakraborty, and Alexander Goesmann. 2020. “Platon: Identification and Characterization of Bacterial Plasmid Contigs in Short-Read Draft Assemblies Exploiting Protein Sequence-Based Replicon Distribution Scores.” Microbial Genomics 6 (10). 10.1099/mgen.0.000398.

31. Seemann, Torsten. 2014. “Prokka: Rapid Prokaryotic Genome Annotation.” Bioinformatics 30 (14): 2068–69.

32. Tiedje, James M., Fang Wang, Célia M. Manaia, Marko Virta, Hongjie Sheng, Liping Ma, Tong Zhang, and Edward Topp. 2019. “Antibiotic Resistance Genes in the Human-Impacted Environment: A One Health Perspective.” Pedosphere 29 (3): 273–82.

33. Vanderhaeghen, Wannes, and Jeroen Dewulf. 2017. “Antimicrobial Use and Resistance in Animals and Human Beings.” The Lancet. Planetary Health 1 (8): e307–8.

34. Wind, Lauren, Ishi Keenum, Suraj Gupta, Partha Ray, Katharine Knowlton, Monica Ponder, W. Cully Hession, Amy Pruden, and Leigh-Anne Krometis. 2021. “Integrated Metagenomic Assessment of Multiple Pre-Harvest Control Points on Lettuce Resistomes at Field-Scale.” Frontiers in Microbiology 12 (July): 683410.

35. Wood, Derrick E., and Steven L. Salzberg. 2014. “Kraken: Ultrafast Metagenomic Sequence Classification Using Exact Alignments.” Genome Biology 15 (3): R46.

36. Xiao, Ke-Qing, Bing Li, Liping Ma, Peng Bao, Xue Zhou, Tong Zhang, and Yong-Guan Zhu. 2016. “Metagenomic Profiles of Antibiotic Resistance Genes in Paddy Soils from South China.” FEMS Microbiology Ecology 92 (3). 10.1093/femsec/fiw023.

37. Xia, Yu, An-Dong Li, Yu Deng, Xiao-Tao Jiang, Li-Guan Li, and Tong Zhang. 2017. “MinION Nanopore Sequencing Enables Correlation between Resistome Phenotype and Genotype of Coliform Bacteria in Municipal Sewage.” Frontiers in Microbiology 8 (October): 2105.

38. Xiong, Wenguang, Mei Wang, Jinjun Dai, Yongxue Sun, and Zhenling Zeng. 2018. “Application of Manure Containing Tetracyclines Slowed down the Dissipation of Tet Resistance Genes and Caused Changes in the Composition of Soil Bacteria.” Ecotoxicology and Environmental Safety 147 (January): 455–60.

39. Xu, Han, Zeyou Chen, Ruiyang Huang, Yuxiao Cui, Qiang Li, Yanhui Zhao, Xiaolong Wang, Daqing Mao, Yi Luo, and Hongqiang Ren. 2021. “Antibiotic Resistance Gene-Carrying Plasmid Spreads into the Plant Endophytic Bacteria Using Soil Bacteria as Carriers.” Environmental Science & Technology 55 (15): 10462–70.

40. Yang, Lu, Wuxing Liu, Dong Zhu, Jinyu Hou, Tingting Ma, Longhua Wu, Yongguan Zhu, and Peter Christie. 2018. “Application of Biosolids Drives the Diversity of Antibiotic Resistance Genes in Soil and Lettuce at Harvest.” Soil Biology and Biochemistry. 10.1016/j.soilbio.2018.04.017.

41. Yin, Xiaole, Xiao-Tao Jiang, Benli Chai, Liguan Li, Ying Yang, James R. Cole, James M. Tiedje, and Tong Zhang. 2018. “ARGs-OAP v2.0 with an Expanded SARG Database and Hidden Markov Models for Enhancement Characterization and Quantification of Antibiotic Resistance Genes in Environmental Metagenomes.” Bioinformatics. 10.1093/bioinformatics/bty053.

42. Zhang, Yu-Jing, Hang-Wei Hu, Qing-Lin Chen, Brajesh K. Singh, Hui Yan, Deli Chen, and Ji-Zheng He. 2019. “Transfer of Antibiotic Resistance from Manure-Amended Soils to Vegetable Microbiomes.” Environment International 130 (September): 104912.

43. Zhang, Yu-Jing, Hang-Wei Hu, Qing-Lin Chen, Hui Yan, Jun-Tao Wang, Deli Chen, and Ji-Zheng He. 2019. “Manure Application Did Not Enrich Antibiotic Resistance Genes in Root Endophytic Bacterial Microbiota of Cherry Radish Plants.” Applied and Environmental Microbiology. 10.1128/aem.02106-19.

44. Zhang, Yu-Jing, Hang-Wei Hu, Min Gou, Jun-Tao Wang, Deli Chen, and Ji-Zheng He. 2017. “Temporal Succession of Soil Antibiotic Resistance Genes Following Application of Swine, Cattle and Poultry Manures Spiked with or without Antibiotics.” Environmental Pollution. 10.1016/j.envpol.2017.09.074.

45. Zhao, Renxin, Jie Feng, Xiaole Yin, Jie Liu, Wenjie Fu, Thomas U. Berendonk, Tong Zhang, Xiaoyan Li, and Bing Li. 2018. “Antibiotic Resistome in Landfill Leachate from Different Cities of China Deciphered by Metagenomic Analysis.” Water Research 134 (May): 126–39.

46. Zhu, Bokai, Qinglin Chen, Songcan Chen, and Yong-Guan Zhu. 2017. “Does Organically Produced Lettuce Harbor Higher Abundance of Antibiotic Resistance Genes than Conventionally Produced?” Environment International 98 (January): 152–59.

47. Zhu, Yong-Guan, Timothy A. Johnson, Jian-Qiang Su, Min Qiao, Guang-Xia Guo, Robert D. Stedtfeld, Syed A. Hashsham, and James M. Tiedje. 2013. “Diverse and Abundant Antibiotic Resistance Genes in Chinese Swine Farms.” Proceedings of the National Academy of Sciences of the United States of America 110 (9): 3435–40.

