## supplementary information for "Metagenomic Analysis Reveals the Transfer of Antibiotic Resistance from Manure-Fertilized Soil to Carrot Microbiome"

### DNA extraction and Hi-seq Sequencing Methods

#### Extraction of DNA from soil, carrot peel wash and soil wash

To obtain soil DNA, 250 mg of soil was extracted with the DNeasy PowerSoil kit (Qiagen, Canada) following the manufacturer's instructions. The final elution volume was 100 µL. To extract DNA from carrot peel wash and soil wash, 250 mg of wash pellets were used for extraction using the DNeasy PowerSoil kit (Qiagen, Canada) following the manufacturer's instructions with a final elution volume of 100 µL. The DNA concentrations and quality were determined using a NanoDrop ND1000 microspectrophotometer (NanoDrop Technologies, Wilmington, DE).

#### Next generation sequencing protocol

Two Carrot peel wash and two soil wash DNA samples from vegetables grown in ground without application of manure, or with swine manure were chosen for next generation sequencing. For each sample, 800 ng of DNA, with an OD260/280 between 1.8 and 2.0 were sent to The Centre for Applied Genomics (The Hospital for Sick Children, Toronto, ON) for Nextera XT library preparation and Illumina high-throughput sequencing (HTS). HTS was performed using the paired-end sequencing strategy (PE150 bp) on HiSeq 4000 platform.

### Metagenomic Reads information

After QC, there was an average of 111,516,766 sequences in the natural carrot peel sample, 99,591,260 in the manured peel sample, while in the soil sample, control group has 97,448,687 reads, and 107,008,589 reads were sequenced in the manured group. It was clear that rarefaction curves reached their asymptotes consistently, indicating that read depth was likely sufficient for ARGs and bacterial communities (Figure S1). 73.29% of the post-QC reads were assembled into 92,036 contigs with N50 of 1,960bp containing 173,447 ORFs on the carrot peel. In the soil sample, 69.47% reads were assembled into 74,483 contigs with N50 of 1,920bp containing 156,542 ORFs. Based on HMM-based function prediction, we annotated 3,659 and 3,771

ORFs as ARG-like ORFs of carrot peel and soil samples, respectively, which were located in 92,036 and 74,483 contigs individually.

Table S1 Statistics on the metagenomic datasets and metagenomic assembly applied in this study.

| | Data size, Gb | Assembly efficiency,% | ARGs-contigs No. |
|---|---|---|---|
| Peel-control | 78 | 73.29% | 92036 |
| Peel-treated | 70 | 73.29% | 92036 |
| Soil-control | 68 | 69.47% | 74483 |
| Soil-treated | 74 | 69.47% | 74483 |

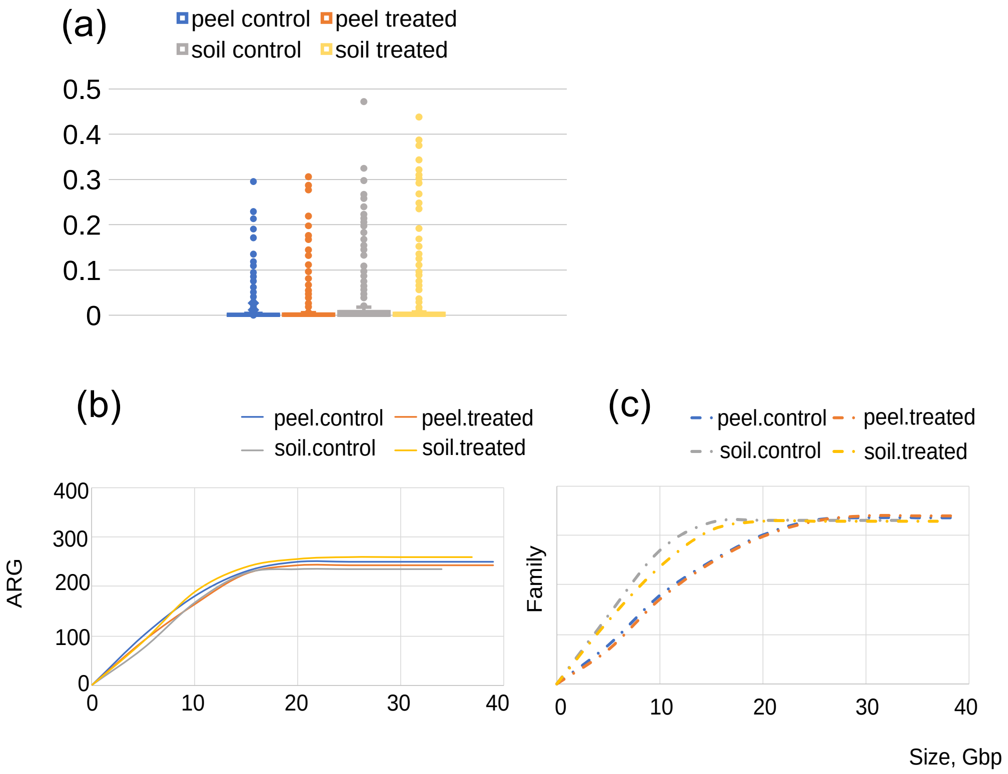


Figure S1 (a) The boxplots show the comparison of ARGs abundance among all soil and peel groups (Paired-sample *t-test*). (b) rarefaction analysis of ARG types along with the increase of the data size of all samples (c) rarefaction analysis of family types along with the increase of the data size of all samples

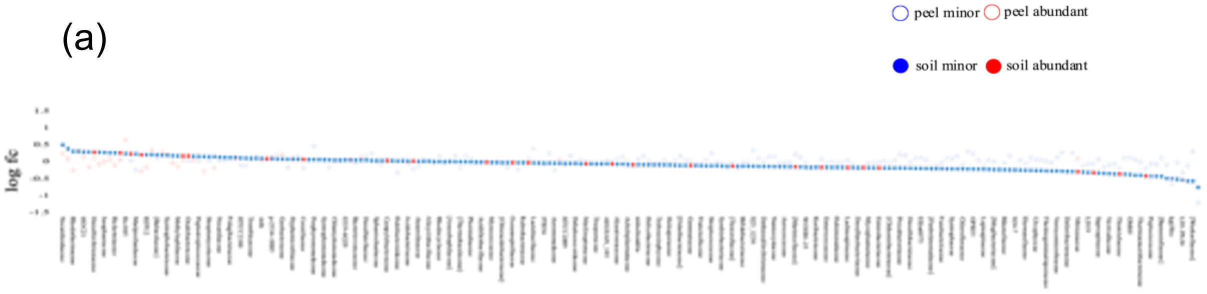

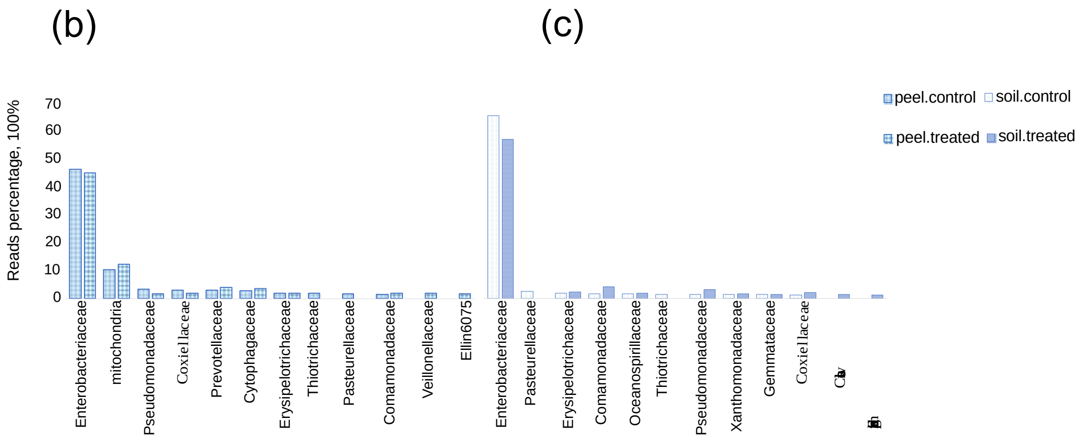


Figure S2 (a) the overall trend of the lograithm of the fold change of the abundance of family OTU on the carrot and soil based on the metagenomic reads (b) The most 10 dominating families of natural peel and manured peel sample (c) The most 10 dominating families of natural soil and manured soil sample

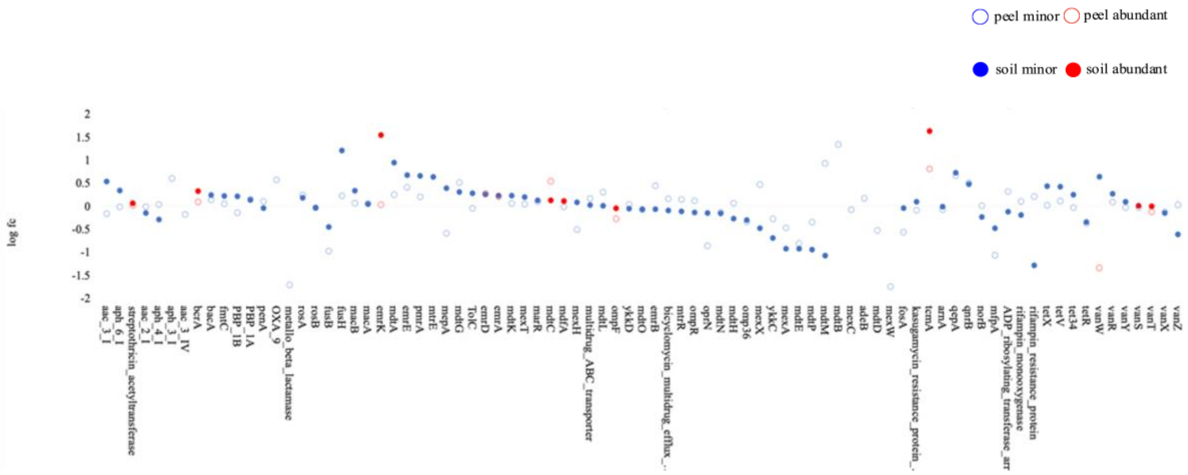


Figure S3 The overall trend of the lograithm of the fold change of the abundance of ARGs on the carrot and soil based on the metagenomic assembly

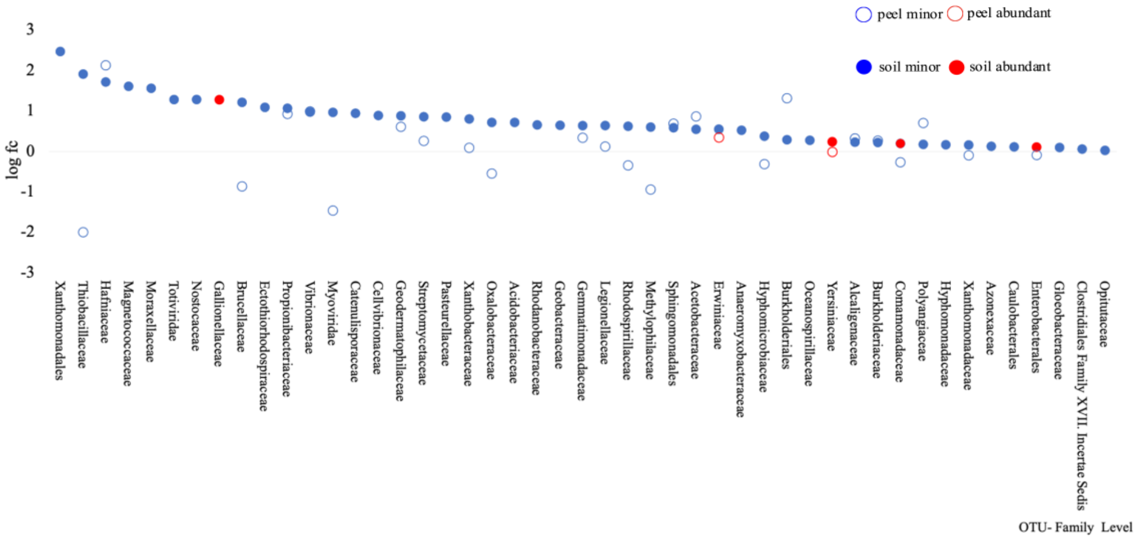


Figure S4 The overall trend of the lograithm of the fold change of the abundance of family OUT on the carrot and soil based on the metagenomic assembly

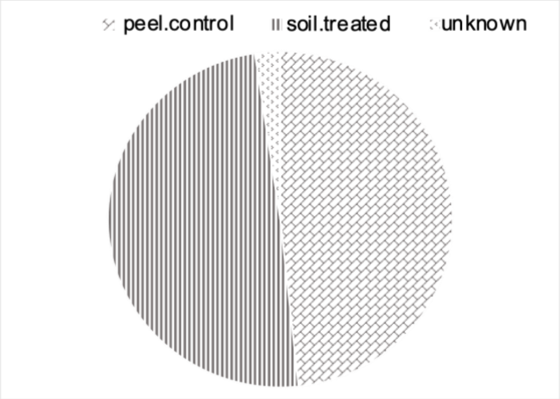


Figure S5 Source tracking result of ARGs based on the metagenomic reads, which regards manured peel as the sink, natural peel, manured soil as the source

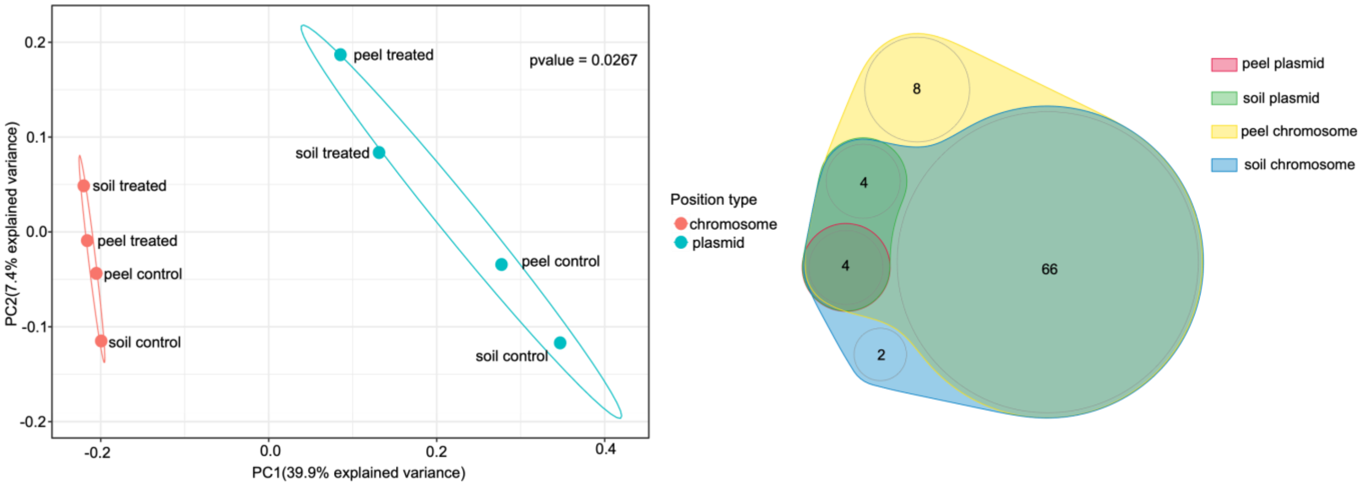


Figure S6 (a) PCoA analysis of ARGs mediated by plasmid and chromosome on the peel and in the soil sampes (b) Venn analysis of ARG types mediated by plasmid and chromosome on the peel and in the soil samples
